# Predictive Modeling of Cancer Cell Growth Kinetics with Machine Learning

**DOI:** 10.64898/2026.09.19.752932

**Authors:** Kaiqiang Hu, Yunyang Zhang, Li Feng, Qiuyuan Yang, Zhe Li, Pengwei Pan, Fang He

**Author notes:** **Correspondence to:** Fang He, PhD. Kaiqiang Hu and Yunyang Zhang contributed equally to this work.

## Abstract

Quantitative characterization of cell proliferation is central to preclinical drug discovery. Here, we evaluated Random Forest (RF) regression for predicting confluence-based cell growth trends using data from human cancer cell lines and benchmarked its performance against the widely used logistic and Gompertz models. The RF model achieved higher predictive accuracy within the evaluated dataset, particularly at higher initial seeding densities. We further extended the RF framework to predict growth curves across seeding densities. Collectively, these findings highlight the potential of data-driven modeling to complement conventional mathematical approaches and support more informed cell-culture design.

## 1. Introduction

Cell culture is a cornerstone of in vitro biomedical research [1]. Cell growth curves provide a quantitative representation of proliferation dynamics and are widely used to assess how pharmacological agents and experimental conditions influence cell behavior [2]. Because proliferation unfolds continuously over time, early measurements may contain information about subsequent growth. However, biological heterogeneity and measurement variability can obscure these temporal patterns. Modeling the growth process may therefore reduce the need for frequent sampling while improving the efficiency and reproducibility of longitudinal growth assessment.

Cell proliferation commonly progresses through lag, exponential-growth, and stationary phases, which together produce an approximately sigmoidal trajectory [3]. This behavior has motivated the widespread use of exponential, logistic, and Gompertz models to describe population growth [4]. Although these models offer interpretable summaries of growth kinetics, their predictive performance may be limited when genetic, environmental, and intercellular factors introduce dynamics not represented by their underlying assumptions [5,6]. More flexible formulations have therefore been developed. For example, Roy et al. [7] extended the logistic model within a stochastic framework to incorporate density regulation, stochastic variation, cooperative behavior, and negative feedback. Nevertheless, models optimized for a restricted set of conditions may not generalize readily to other experimental settings [8,9], while additional terms introduced to represent processes such as adaptation-related lag can substantially increase model complexity [10,11].

Advances in predictive biology have established data-driven modeling as a complementary strategy for analyzing complex biological systems [12]. Machine-learning methods can capture nonlinear associations between multiple inputs and outcomes without requiring every interaction to be specified a priori [13,14]. Koyama et al. [15], for example, applied machine learning to predict the growth of Escherichia coli O157 across temperature conditions. Building on this broader predictive paradigm, we investigated whether a machine-learning framework could model growth trajectories across human cancer cell lines and initial seeding densities.

This study aimed to develop and evaluate an RF regression model for predicting cell-growth trends across multiple cancer cell lines and initial seeding densities. We benchmarked its predictive performance against logistic and Gompertz models and examined how the temporal input measurements and initial seeding density were associated with model performance. We also assessed whether RF regression could predict growth trajectories across seeding densities.

## 2. Materials and Methods

### 2.1. Cell culture

Cells were revived and cultured overnight in T75 flasks before growth-curve analysis. The next day, cell density was measured, and cell suspensions were prepared according to the experimental design and seeded into 384-well plates. Cells were maintained in a humidified incubator at 37 °C and 5% CO2. Whole-well confluence was recorded from day 0 through day 7 using an IncuCyte system.

### 2.2. Development of the machine-learning framework

#### 2.2.1. Constructing mathematical models

Prediction analyses initially employed the logistic and Gompertz models. In these models, the change in cell population N over time t is described by Equations (1) and (2), respectively:

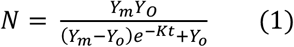

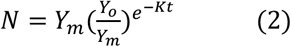

where Y0, Ym, and K represent the minimum cell population, maximum cell population, and rate constant, respectively.

#### 2.2.2. Constructing machine learning models

Cell-growth observations were randomly partitioned into training and test sets, with 70% allocated to model training and 30% reserved for evaluation. An RF regression model comprising 500 decision trees was fitted to the training data. Model implementation was performed in Python using the scikit-learn library.

#### 2.2.3. Error metrics for evaluating models

The coefficient of determination (R^2^) was used to quantify the proportion of variance explained by the model [16]. Mean relative error (MRE) was used to characterize the discrepancy between predicted and observed values [17]. Higher R^2^ values and lower MRE values indicate better predictive performance, as defined in Equations (3) and (4):

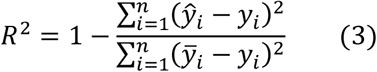

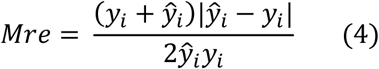

where ŷyi, and 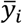 are the predicted, actual, and mean values, respectively.

### 2.3. Model analysis

#### 2.3.1. Analysis of input characteristic variables

Pearson’s correlation coefficient (PCC) was used to quantify the strength and direction of pairwise linear associations among the continuous input variables [18]. PCC was calculated as shown in Equation (5):

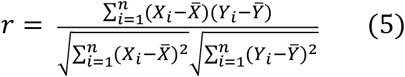

where r denotes the PCC between two variables; Xi and Yi are the corresponding observed values; and the barred X and Y terms denote their respective means. Values of r range from -1 to 1, with values approaching 1 or -1 indicating strong positive or negative linear correlations, respectively, and a value of 0 indicating no linear correlation [19].

Feature importance was subsequently evaluated for all input variables. During RF model construction, the contribution of each feature was quantified from the error reduction associated with splits using that feature across the decision trees. These contributions were summed across trees and normalized to derive the model-specific feature-importance scores [20]. To further assess the contribution of different input combinations, additional models were constructed using grouped feature sets and evaluated by five-fold cross-validation.

#### 2.3.2. Analysis of initial seeding density

Samples were stratified into seven groups according to initial seeding density (50, 100, 200, 400, 800, 1,600, and 3,200 cells/well). We determined the number of samples at each density in the training and test sets and counted test samples with MRE values below 0.1 or above 0.4. The proportions of samples in these MRE categories were then calculated for each seeding-density group.

The association between sample MRE and cell-line doubling time (Td) was then examined. For test samples seeded at 50 cells/well, the 50 samples with the lowest MRE and the 50 samples with the highest MRE were assigned to lower-error and higher-error groups, respectively. The same procedure was applied to samples seeded at 100 cells/well using the 45 samples with the lowest and highest MRE values. Cell-line Td was calculated using Equation (6):

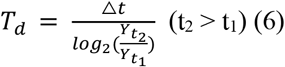

where Yt1 and Yt2 are the cell-confluence values at times t1 and t2, respectively, and Δt is the interval between t2 and t1.

The calculated Td values were summarized for each group. Wilcoxon rank-sum tests were used to evaluate differences between the lower-error and higher-error groups at the same initial seeding density.

## 3. Results

### 3.1. Model performance evaluation

The RF model achieved an R^2^ of 0.957 on the test data (Table 1). For descriptive comparison, model performance was categorized according to the MRE between the observed and predicted growth curves. Predictions with MRE below 0.1 closely matched the observed curves, whereas values of 0.1-0.2 indicated a general correspondence. MRE values above 0.2 reflected progressively larger deviations, and values above 0.4 indicated marked divergence between predicted and observed curves (Figure 1). Overall, 865 samples had an MRE below 0.2, representing nearly 75% of the test set and demonstrating the predictive performance of the RF model within the evaluated dataset.

**Table 1.** R^2^ and sample MRE distributions for the different models in the test set.

| Model | $R^2$ of testing set | Testing samples | | | |
| --- | --- | --- | --- | --- | --- |
| | | $MRE \in [0,0.1]$ | $MRE \in (0.1,0.2]$ | $MRE \in (0.2,0.4]$ | $MRE \in (0.4,+\infty)$ |
| RF | 0.957 | 627 | 238 | 206 | 98 |
| Logistic | -6.708 | 267 | 227 | 277 | 398 |
| Gompertz | -7.791 | 226 | 224 | 290 | 429 |

**Figure 1.**
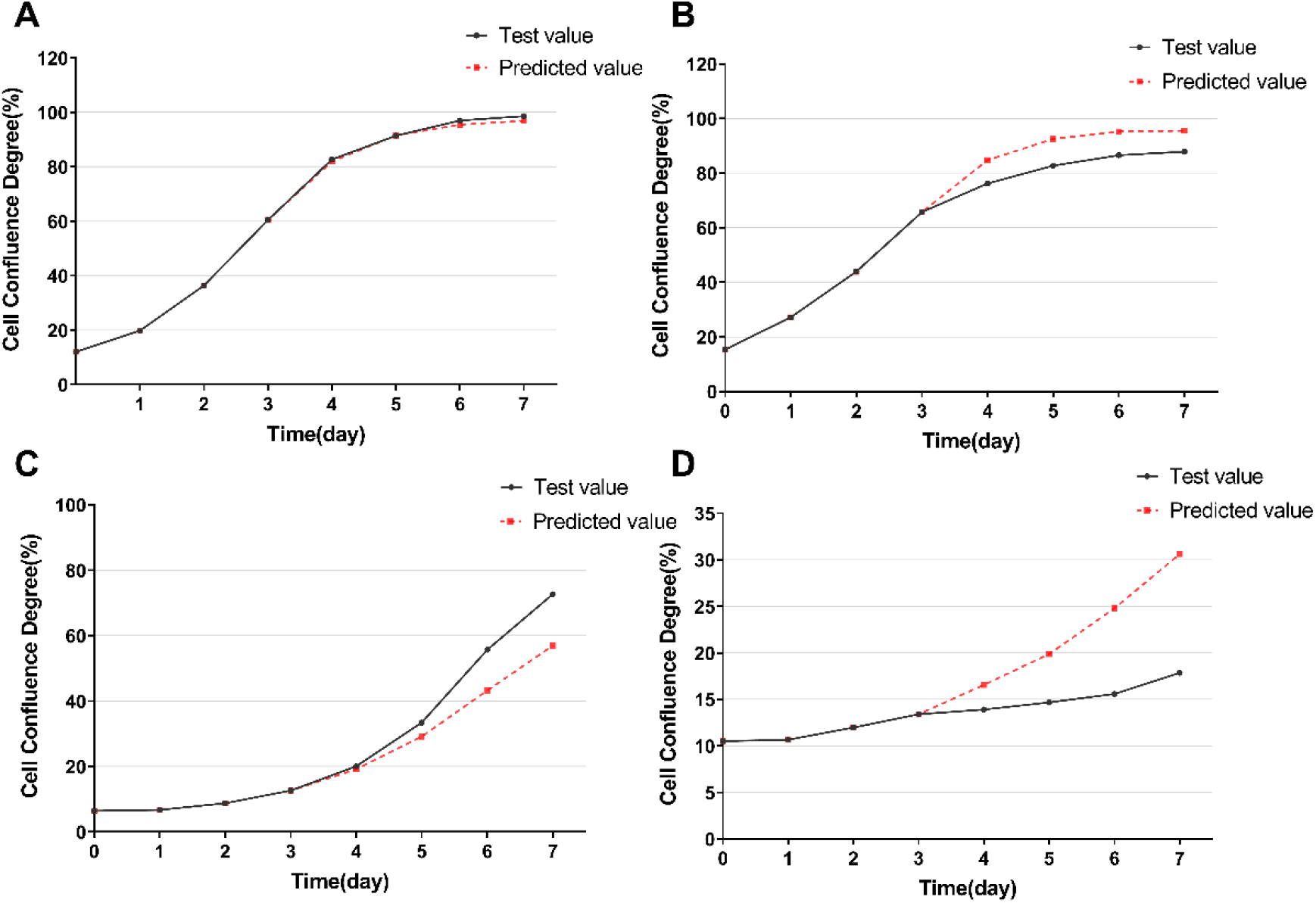
Representative comparisons between predicted and observed curves at relative errors of 0.01 (**A**), 0.1 (**B**), 0.2 (**C**), and 0.4 (**D**).

### 3.2. The RF model outperforms the mathematical models in predictive accuracy

We compared the predictive performance of the RF, logistic, and Gompertz models using the metrics reported in Table 1. MRE values below 0.1 were observed for 53.6% of RF predictions, compared with 19.3% of Gompertz-model predictions and 22.8% of logistic-model predictions. Conversely, 57.7% of logistic-model predictions and 61.5% of Gompertz-model predictions had MRE values above 0.2 (Figure 2A). In the test set, the RF model had a lower MRE (0.15293) than the logistic (0.40233) and Gompertz (0.43547) models (Figure 2B). The error distributions of the two mathematical models were also broader and contained more high-MRE observations than that of the RF model (Figure 2C). Figure 2D illustrates representative cases in which the mathematical-model fits converged prematurely or failed to converge.

**Figure 2.**
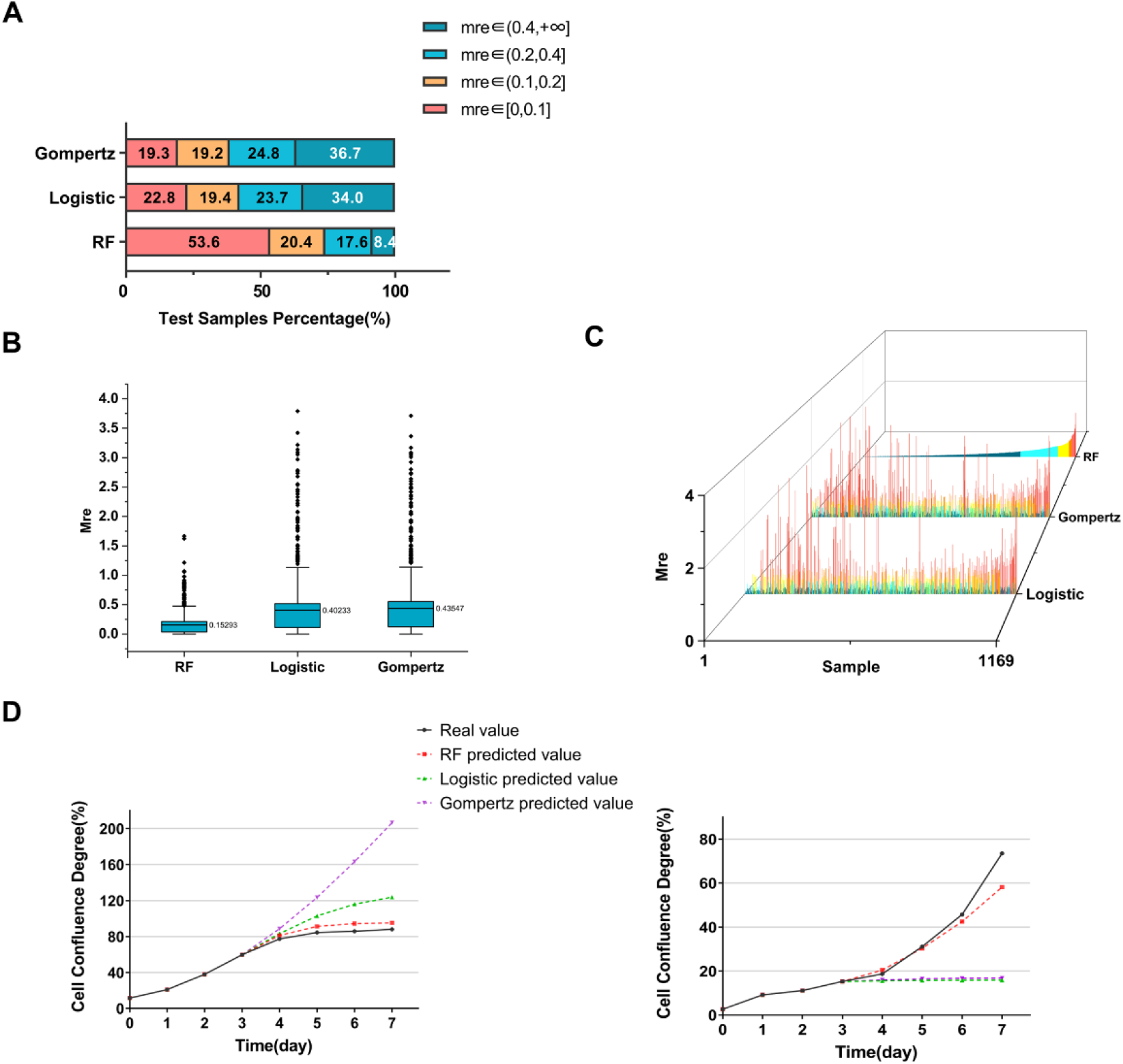
Performance comparison of the RF and mathematical models. **A**. Distribution of MRE across error ranges for the three models. **B**. MRE distributions for the different models. **C**. Sample-level MRE for the different models. **D**. Representative errors in mathematical-model predictions of cell growth.

### 3.3. Day-3 measurements dominate RF feature importance

To identify inputs associated with model predictions, we calculated pairwise correlations among cell-confluence measurements from day 0 through day 3. All correlation coefficients exceeded 0.7, indicating strong correlations among the input features (Figure 3A). In the RF model, the day-3 measurement had a feature-importance score of 0.9164 and was therefore the dominant predictor by the model’s split-based importance measure (Figure 3B and Table S1). Comparisons of R^2^ across models with different input combinations showed that predictive performance was highest when all four days of growth data were included. Models that included day-3 measurements also tended to perform better (Figure 3C).

**Figure 3.**
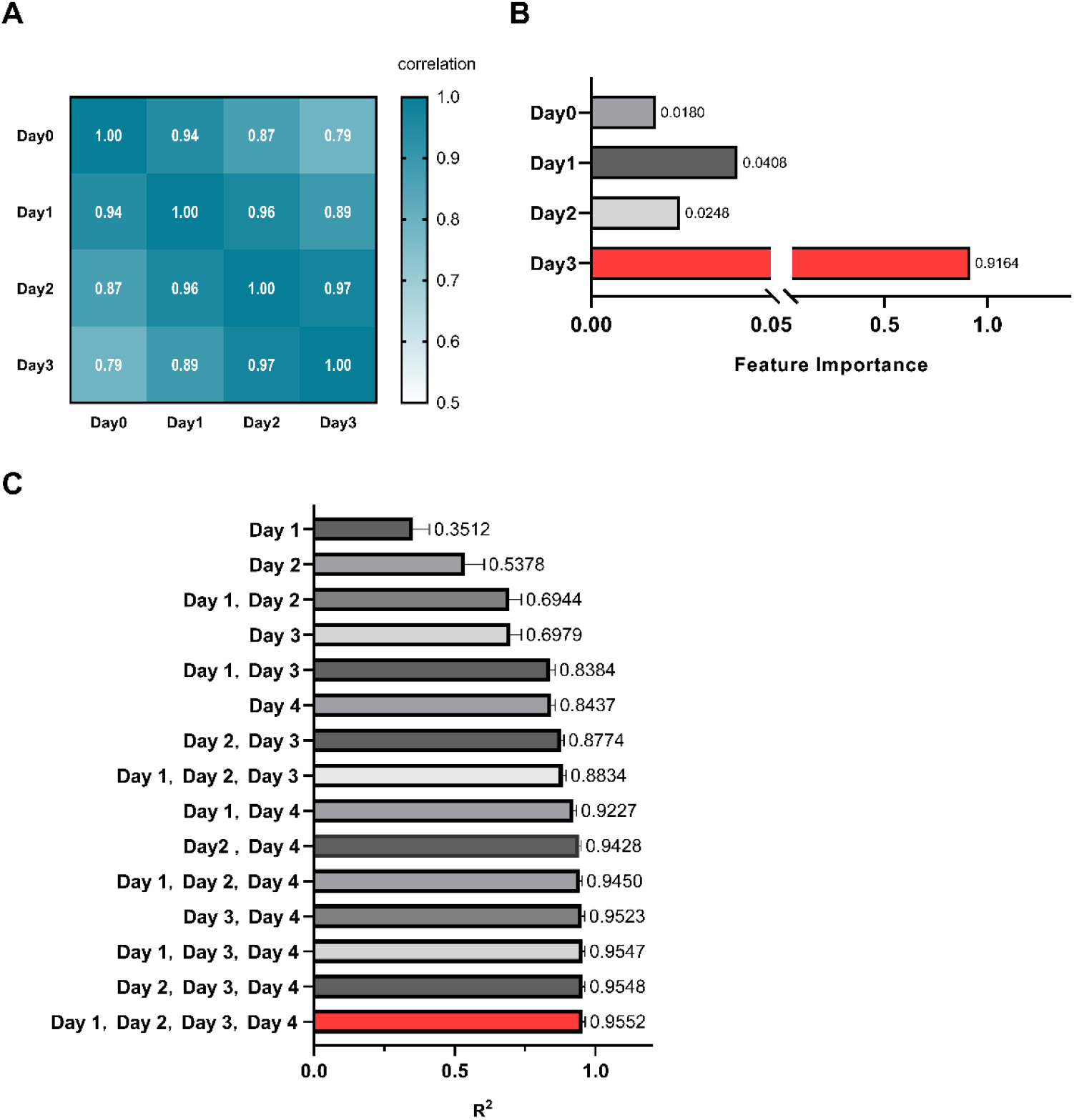
Contributions of input features to model performance. **A**. Correlation matrix of the input features. Correlations become stronger as the coefficient approaches 1. **B**. Feature-importance scores in the RF model. **C**. Performance of models grouped by input-feature combinations.

### 3.4. Predictive performance varies with initial seeding density

To investigate factors associated with predictive performance, we compared MRE across initial seeding-density groups (Table S2). Samples with MRE below 0.1 were predominantly seeded at 800 cells/well or higher, whereas samples with MRE above 0.4 were concentrated at 200 cells/well or lower (Figure 4A). MRE generally decreased as initial seeding density increased (Figure 4B), indicating lower predictive performance at lower densities in this dataset. Samples seeded below 200 cells/well also exhibited higher coefficients of variation (CVs), consistent with greater relative variability at low seeding densities (Figure 4C).

**Figure 4.**
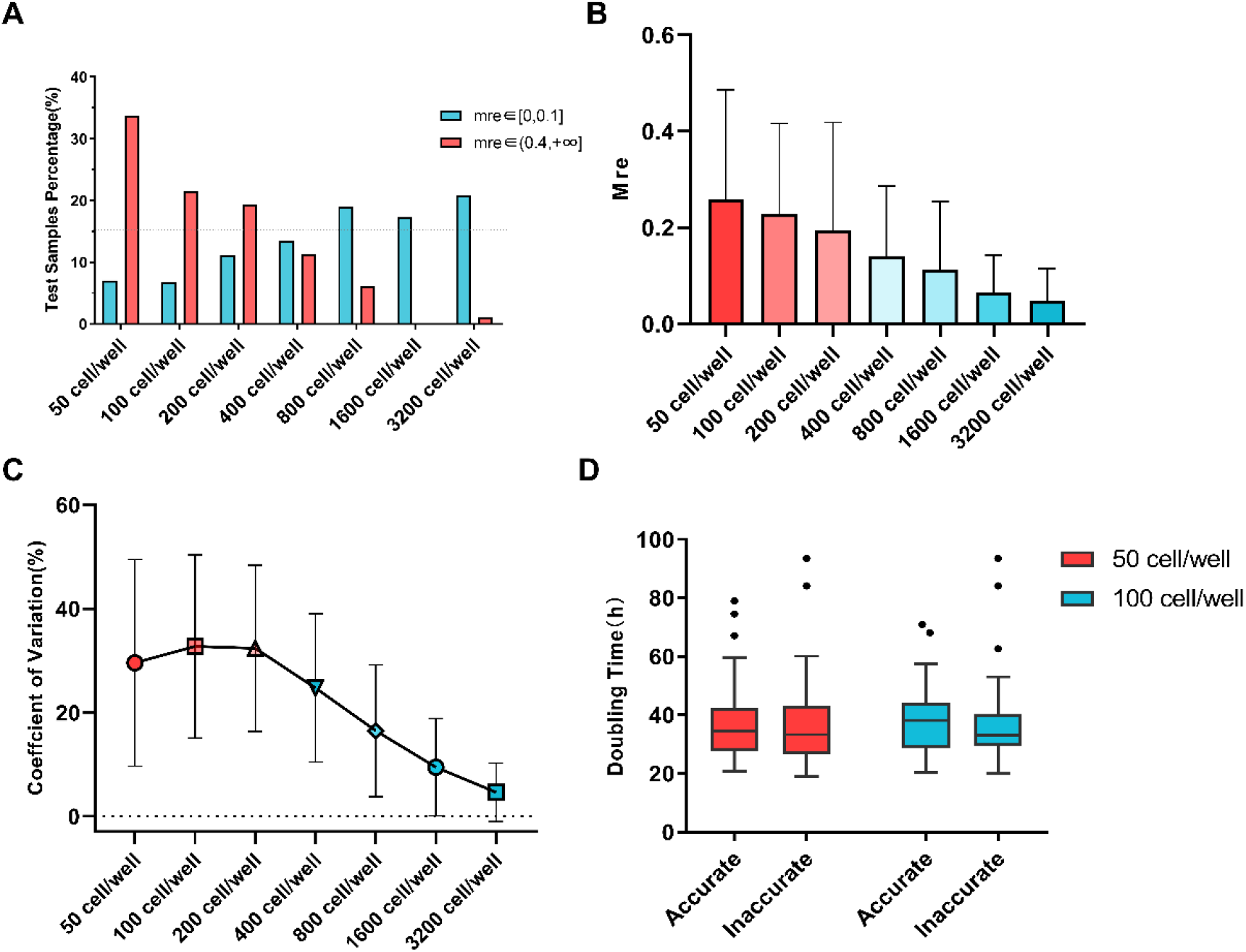
Association between initial seeding density and model performance. **A**. MRE distributions across initial seeding densities. **B**. MRE by initial seeding density. **C**. Coefficient of variation (CV) in cell confluence from day 4 through day 7 across initial seeding densities. **D**. Distribution of cell-line doubling time (Td) by prediction-error group at low seeding densities.

We next examined whether Td differed between the lower-error and higher-error groups at low initial seeding densities. The Td distributions were similar between groups (Figure 4D and Table 2). Wilcoxon rank-sum tests yielded P = 0.231 for samples seeded at 50 cells/well and P = 0.372 for samples seeded at 100 cells/well. Both P values exceeded 0.05; thus, neither comparison detected a statistically significant difference in Td between the two error groups.

**Table 2.**
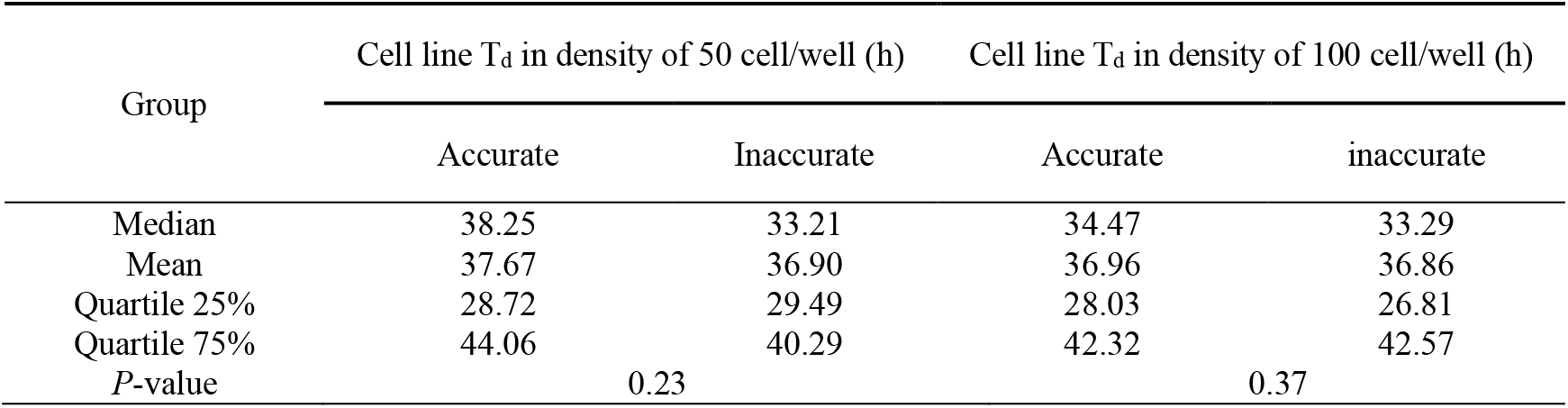
Statistics of cell line T^d^ at low initial cell densities

| Group | Cell line T<sub>d</sub> in density of 50 cell/well (h) |  | Cell line T<sub>d</sub> in density of 100 cell/well (h) |  |
| --- | --- | --- | --- | --- |
|  | Accurate | Inaccurate | Accurate | inaccurate |
| Median | 38.25 | 33.21 | 34.47 | 33.29 |
| Mean | 37.67 | 36.90 | 36.96 | 36.86 |
| Quartile 25% | 28.72 | 29.49 | 28.03 | 26.81 |
| Quartile 75% | 44.06 | 40.29 | 42.32 | 42.57 |
| *P*-value | 0.23 |  | 0.37 |  |

### 3.5. Prediction of cell-growth trends across seeding densities

In the test set for the density model, 79% of samples had an MRE below 0.08, and predictions within this range generally followed the observed curves (Figure 5A and 5B). Joint prediction of the growth curves at 1,600 and 800 cells/well produced better performance than predicting each curve separately (Figure 5B and Table S3). Feature-importance analysis assigned relatively high importance to mid-phase measurements at 3,200 cells/well and late-phase measurements at 400 cells/well (Figure 5C and Table S4). These model-specific patterns may reflect differences in the information captured at distinct growth phases and seeding densities.

**Figure 5.**
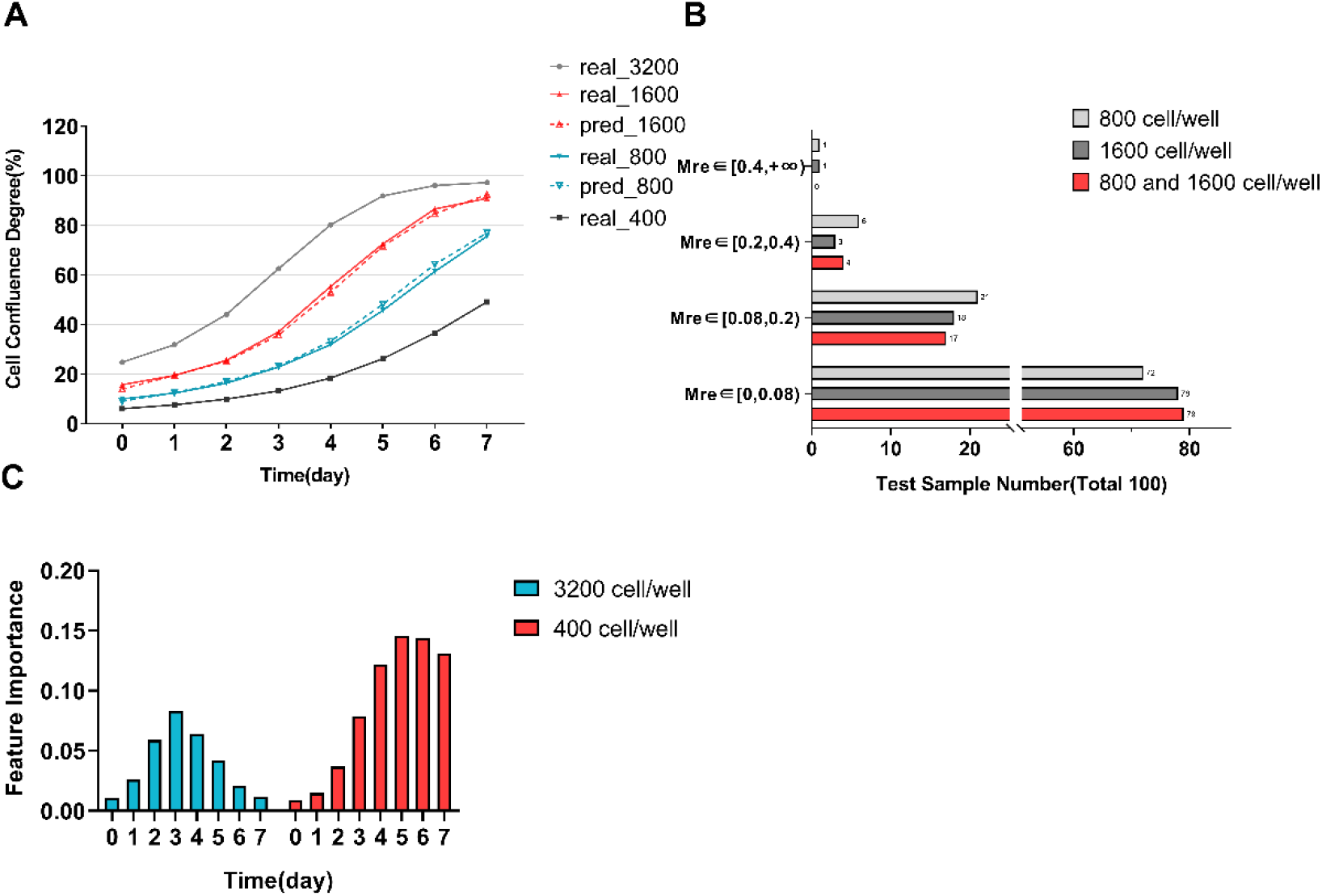
Evaluation of the density-based cell-growth prediction model. **A**. Representative prediction from the density model. **B**. MRE distributions for models constructed with different outputs. **C**. Feature-importance scores in the density model.

## 4. Discussion

In this study, we developed an RF regression framework to predict confluence-based cell-growth trajectories. Cellular responses are inherently variable and can exhibit distinct nonlinear dynamics across cell types [21]. Conventional mathematical models remain valuable for characterizing population growth, but their predictive performance can decline when observed trajectories depart from their parametric assumptions. Indeed, several cell lines in our dataset deviated from the canonical sigmoidal profile at specific seeding densities, limiting the ability of the logistic and Gompertz models evaluated here to capture their growth behavior [22].

RF regression is well suited to nonlinear and multi-output prediction and can be robust to certain forms of noise through random sampling and feature selection [23]. In the present test dataset, the RF model produced lower prediction errors than the logistic and Gompertz models (Figure 2A). Its performance nevertheless varied with initial seeding density, with higher errors at lower densities and improved performance at higher densities. Cell-to-cell heterogeneity may contribute more strongly to population-level variability when relatively few cells are initially present [24]. Using Monte Carlo simulations, Koutsoumanis and Lianou showed that variability in bacterial population growth decreased as the initial cell number increased and became negligible above 100 cells [25]. Although bacterial findings may not translate directly to cancer-cell cultures, they provide a plausible conceptual framework for the density-associated pattern observed here: predictive performance improved substantially in wells seeded with more than 200 cells (Figure 4A and 4B).

Variation in cell-line doubling time was considered as one potential explanation for the lower predictive performance observed at seeding densities of 50 and 100 cells/well. However, no statistically significant difference in doubling time was detected between the lower-error and higher-error groups (Figure 4D). Doubling time reflects intrinsic cell-line characteristics but is also shaped by environmental conditions. Cells can approach their maximum growth rate when space and nutrients are abundant and slow as population density increases [26]. We therefore hypothesize that cells seeded at low density undergo more variable growth dynamics during the first 4 days, making their subsequent trajectories more difficult to predict. This interpretation remains to be tested directly.

The density model further demonstrated that RF regression could represent nonlinear temporal dynamics while capturing information shared across curves generated at different seeding densities. Notably, joint prediction of the two intermediate-density curves outperformed separate prediction of each curve (Figure 5B), suggesting that the shared output structure contained information that improved overall prediction. Feature-importance analysis highlighted mid-phase measurements at high initial density and late-phase measurements at low initial density (Figure 5C). During these intervals, confluence changed across a broader dynamic range, potentially increasing the predictive information provided by these measurements.

In summary, we developed an RF regression model for predicting cancer-cell growth trajectories and benchmarked it against logistic and Gompertz models. Within the evaluated dataset, the RF model produced lower prediction errors, and its performance varied with initial seeding density. The group comparisons did not detect a statistically significant difference in doubling time between lower-error and higher-error samples at low seeding densities. We also demonstrated the use of RF regression to predict growth trends across seeding densities. These findings establish a foundation for further evaluation of data-driven cell-growth prediction using independent datasets and broader experimental conditions.

## Glossary

Random Forest (RF): An ensemble machine learning algorithm for classification or regression
Pearson%s Correlation Coefficient (PCC): A statistical measure of the linear relationship between two variables
Coefficient of Determination (R^2^): A statistical metric that measures the accuracy of a regression model
Mean Relative Error (MRE): A measure of prediction accuracy, expressed as the average error relative to actual values
Coefficient of Variation (CV): A standardized measure of dispersion, indicating the relative variability of data points
Doubling Time (T_d_): The period needed for a cell population to double its number
Logistic model: A mathematical function that describes an S-shaped curve
Gompertz model: A mathematical function that describes an S-shaped curve with a late peak
Five-Fold Cross-Validation: A method of model validation that involves dividing the data into five equal parts
Wilcoxon test: A non-parametric statistical test that compares the medians of two samples.

## Data availability

The data supporting the findings of this study are available from the corresponding author upon reasonable request. The data are not publicly available because of privacy or ethical restrictions. The relevant code is available for download from the following GitHub repository: https://github.com/Unicorn0512/Cell_Confluence_Prediction

## Author contribution

**Kaiqiang Hu:** Conceptualization, Formal analysis, Writing - Original Draft, Writing - Review & Editing. **Yunyang Zhang:** Visualization, Methodology, Writing - Original Draft, Validation. **Li Feng:** Investigation. **Qiuyuan Yang:** Writing - Review & Editing. **Zhe Li**: Writing - Review & Editing. **Pengwei Pan:** Supervision, Conceptualization, Project administration. **Fang He:** Supervision, Funding acquisition, Validation.

## Acknowledgements

We thank the staff who contributed to cell culture and Dr. Zhaozhao Wu for guidance on machine-learning methods. This work was funded by Pharmaron’s internal research and development project Bio-PHAR-YFG-2603003.

